# Frequent intergenotypic recombination between the non-structural and structural genes is a major driver of epidemiological fitness in caliciviruses

**DOI:** 10.1101/2021.02.17.431744

**Authors:** Jackie E Mahar, Maria Jenckel, Nina Huang, Elena Smertina, Edward C Holmes, Tanja Strive, Robyn N Hall

## Abstract

The diversity of lagoviruses (*Caliciviridae*) in Australia has increased considerably. By the end of 2017, five variants from three viral genotypes were present in populations of Australian rabbits, while prior to 2014 only two variants were known. To understand the interactions between these lagovirus variants we monitored their geographical distribution and relative incidence over time through a landscape-scale competition study, and from this, revealed potential drivers of epidemiological fitness. Within three years of the arrival of GI.1bP-GI.2 (RHDV2) into Australia, we observed the emergence of two novel recombinant lagovirus variants, GI.4eP-GI.2 (4e-recombinant) in New South Wales and GI.4cP-GI.2 (4c-recombinant) in Victoria. Although both novel recombinants contain the non-structural genes from benign, rabbit-specific, enterotropic viruses, these variants were recovered from the livers of both rabbits and hares that had died acutely. This suggests that determinants of host and tissue tropism for lagoviruses are associated with the structural genes, and that tropism is intricately connected with pathogenicity. Phylogenetic analyses demonstrated that the 4c-recombinant emerged independently on multiple occasions, with five distinct lineages observed. Both new recombinant variants replaced the previous dominant parental RHDV2 in their respective geographical areas, despite sharing an identical or near-identical (i.e., single amino acid change) major capsid protein with the parental virus. This suggests that epidemiological fitness of these recombinants was not driven by antigenic variation in the capsid, implicating the non-structural genes as key drivers of epidemiological fitness. Molecular clock estimates place the GI4.e recombination event in early to mid-2015, while the five GI.4c recombination events occurred from late 2015 through to early 2017. The emergence of at least six viable recombinant variants within a two-year period highlights an unprecedented frequency of these events, detectable only due to intensive surveillance, and demonstrates the importance of recombination in lagovirus evolution.

## Introduction

Caliciviruses are an important group of vertebrate-infecting viruses. They include noroviruses, the major cause of gastroenteritis in humans worldwide [1], and members of the genus *Lagovirus*, which infect rabbits and hares. Some lagoviruses are hepatotropic and cause an acute fulminant viral hepatitis with a case fatality rate exceeding 90%, while others are enterotropic and are thought to be entirely benign [2]. These are referred to as rabbit haemorrhagic disease viruses (RHDVs) and rabbit caliciviruses (RCVs), respectively.

Lagoviruses are hierarchically classified by their major capsid protein (VP60) type, and less frequently by polymerase type, into genogroups (e.g. GI, GII), genotypes (e.g. GI.1, GI.2, GI.4), and variants (e.g. GI.1a, GI.1b, GI.1c) [3]. The first lagoviruses identified, from hares in Europe in the early 1980s, were those of the GII.1 genotype (EBHSV) [4]. Subsequently, mortality events in *Oryctolagus* rabbits in China in 1984 led to the discovery of genotype GI.1 (RHDV1) viruses [5]. With an increasing diversity of GI.1 viruses, this genotype was further subdivided based on the VP60 phylogeny into several variants. In 1997, the first antigenic variants of GI.1 viruses, now classified as GI.1a (RHDVa), were described in Italy [6] and these spread throughout Europe and more distantly. Then, in 2010, the GI.2 (RHDV2) variant first emerged in France [7–9]. RHDV2 rapidly spread globally, triggering epizootics worldwide in wild and domestic lagomorph populations. As surveillance efforts and technologies improved, non-pathogenic GI.3 (RCV-E1) and GI.4 (RCV-A1, RCV-E2) lagoviruses were identified in the 1990s, first in Europe [10–13] and then in Australia [14] and New Zealand [15]. Non-pathogenic hare lagoviruses (GII.2, GII.3, GII.4, GII.5; HaCV) have also been reported more recently, in 2014 in Europe [16–18] and 2019 in Australia [19].

Lagoviruses, like other caliciviruses, are small, non-enveloped viruses containing a monopartite positive-sense single-stranded RNA genome approximately 7.5 kb in length (Figure 1) [2]. Virus particles contain both genomic RNA (gRNA) and 3’ co-terminal sub-genomic RNAs (sgRNA) approximately 2.5 kb in length [20]. In lagoviruses, the non-structural (NS) genes are situated upstream of the major capsid protein, VP60, all of which are encoded as a single large polyprotein (ORF1), while the 3’ terminal ORF2 encodes a minor structural protein that is presumed to be important for genome release during infection [21]. Lagoviruses encode seven NS proteins, including a helicase (2C-like), a viral genome-linked protein (VpG), a protease (Pro), an RNA-dependent RNA polymerase (RdRp), and three proteins of unknown function (p16, p23, p29). Both gRNA and sgRNA are linked to VpG at their 5’ end and are polyadenylated at their 3’ end [22]. There is considerable homology between the 5’ terminal nucleotide sequences of the gRNA and sgRNA (Figure 1) [22], facilitating recombination at the junction between the RdRp and VP60; this region is highly conserved within and between genogroups.

**Figure 1:**
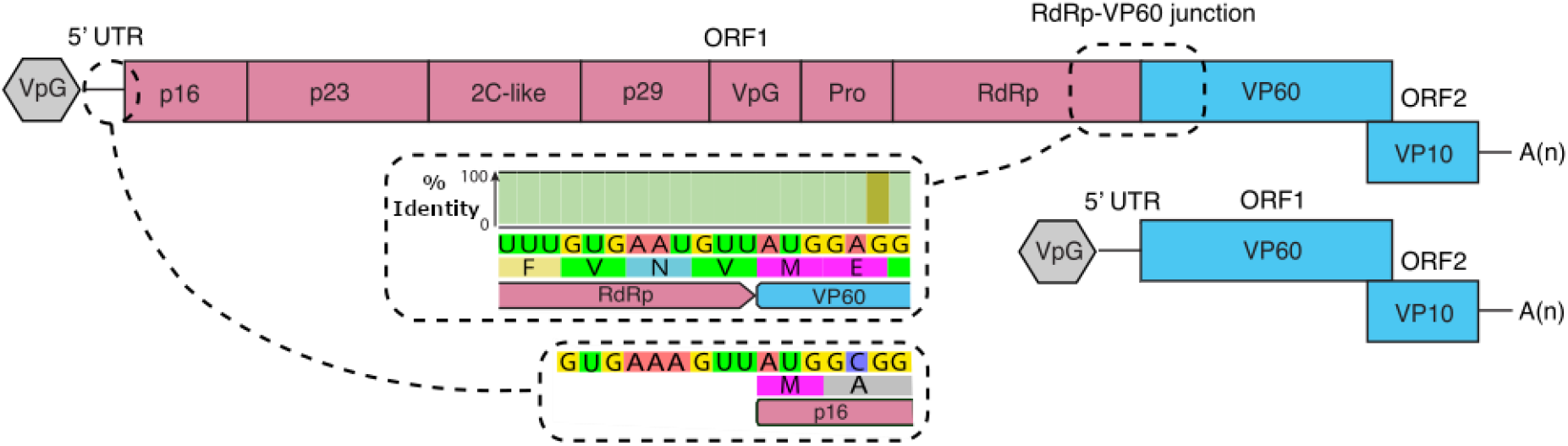
Lagovirus genome organisation. Genomic RNAs (gRNA) encode the non-structural proteins (pink) and structural proteins (blue), while subgenomic RNAs (sgRNA) encode the structural proteins only. Both RNAs are linked to VpG at their 5’ end and are polyadenylated at their 3’ end. Inset boxes demonstrate the nucleotide sequence identity between the 5’ untranslated region (UTR) of the gRNA and the RdRp-VP60 junction. An identity plot (based on n = 478 near-complete GI sequences from GenBank; green indicates 100% identity) is also shown for the junction region.

Recombination is an important evolutionary mechanism in many RNA viruses [23]. In the *Caliciviridae*, recombination frequently occurs at the junction between the NS and structural genes, effectively mixing a set of structural genes with an entirely new set of NS genes [24–26]. Recombinant lagoviruses are defined by the nomenclature [RdRp genotype]P-[capsid genotype], and can include combinations of two pathogenic RHDVs (e.g. GI.1bP-GI.2) as well as the seemingly more common combination of a benign RCV and a pathogenic RHDV (e.g. GI.4eP-GI.1a, GI.4eP-GI.2, GI.3P-GI.2) [24, 27–32]. Retrospective phylogenetic analyses have demonstrated that all GI.2 viruses so far described are recombinants (GI.3P-GI.2), implying that GI.2 is an orphan capsid-type [24]. Intergenogroup recombinants between GI (rabbit and hare) and GII (hare) viruses have also been reported, with two GII.1P-GI.2 viruses recovered from hares in Germany in 2014 and 2019 [33].

Prior to 2016, three lagovirus genotypes (GI.1, GI.2, GI.4) from the G.I genogroup, comprising five distinct variants — GI.4 (RCV-A1), GI.1c (RHDV1), GI.1a (RHDVa-K5), GI.4eP-GI.1a (RHDVa-Aus), GI.1bP-GI.2 (RHDV2) — had been reported in Australia. These variants differ in their host and tissue tropism and in pathogenicity [34]. RCV-A1 is a benign enterotropic virus that has circulated in wild and domestic rabbits, likely since at least the 1950s [14, 35]. RHDV1 and RHDVa-K5 are pathogenic viruses, both deliberately released [36]. RHDVa-Aus and RHDV2 were both exotic incursions, first detected in January 2014 and May 2015, respectively [32, 37]. Phylogenetic analyses suggest that both incursions arose from single point source introductions with subsequent ongoing transmission in Australian rabbit populations [32, 34]. In mid-2017 a novel GI.4eP-GI.2 recombinant virus (4e-recombinant) was detected, comprising the NS sequences of the RHDVa-Aus virus and RHDV2 VP60 sequences [31]. However, the long-term epidemiological significance of this recombinant variant was unclear.

Australia is an ideal setting for understanding the evolution, recombination, and epidemiological interactions of caliciviruses. Australia is an isolated landmass with a large rabbit population that is distributed across most of the country. The genetic diversity of Australian lagoviruses arose from a small number of introduction (or incursion) events, and the parental sequence as well as the timing of the introduction is known in several instances. Since there is little movement of rabbits into the country, the study of lagoviruses can occur in a relatively controlled environment with a defined number of pre-existing pathogenic viruses. Additionally, a robust surveillance system for these viruses has been in place since 2014. The previous lack of diversity of the Australian lagovirus population and sudden diversification of lagovirus variants in 2014/2015 created the perfect natural experimental conditions for monitoring the frequency of successful recombination and drivers of viral evolution. In this study we monitored the epidemiological and evolutionary dynamics of existing lagovirus variants in Australia, while determining the frequency and importance of successful recombination events between these viruses.

## Materials and Methods

### Sample submission

Samples from dead domestic and wild lagomorphs (i.e., rabbits and hares) were submitted either directly or via RabbitScan (https://www.feralscan.org.au/rabbitscan/) to the Commonwealth Scientific and Industrial Research Organisation (CSIRO) by veterinarians, pet owners, landholders, and members of the public as part of ongoing opportunistic lagovirus surveillance. No animal ethics approvals are required for sampling rabbits that are found dead in Australia. Samples were provided either fresh-frozen or stored in an RNA stabilization solution [31]. Additionally, 42 positive samples submitted to the Elizabeth Macarthur Agricultural Institute (EMAI) for RHDV diagnostic testing were forwarded to CSIRO, either as fresh-frozen tissue or as a swab of the cut tissue stored in phosphate buffered gelatin saline (PBGS) [38]. Samples for this study were restricted to those collected between 1 January 2014 (based on the estimated time of incursion of GI.1bP-GI.2 into Australia [34, 39]) and 30 September 2020, and were received from the Australian Capital Territory (ACT, n = 139), New South Wales (NSW, n = 401), the Northern Territory (NT, n = 7), Queensland (QLD, n = 86), South Australia (SA, n = 227), Tasmania (TAS, n = 221), Victoria (VIC, n = 649), and Western Australia (WA, n = 194) (total n = 1,924).

### Initial testing

RNA was extracted from 20 – 30 mg of tissue, predominantly liver or bone marrow, but occasionally spleen, kidney, muscle, ear tip, eyelid, or blowfly maggots retrieved from the carcass. After homogenisation with glass beads using a Precellys 24-dual tissue homogenizer (Bertin Technologies), RNA was extracted using either the Maxwell 16 LEV simplyRNA tissue kit (Promega, Alexandria, NSW) or RNeasy mini kit (Qiagen, Chadstone Centre, VIC) as per manufacturers’ directions. For swab samples, RNA was extracted from 200 µl PBGS using the Purelink viral RNA/DNA mini kit (ThermoFisher Scientific). Samples were screened for known lagoviruses using a broadly-reactive lagovirus SYBR-green based RT-qPCR targeting VP60 and sgRNA, as described previously [31].

### Variant identification

Of the samples that were positive by screening RT-qPCR (n = 1,209), 175 were sequenced previously for other studies [31, 32, 34, 37, 40]. The remaining samples were screened for recombination by RT-PCR and sequencing. RT-PCR primers were designed that spanned the RdRp-VP60 junction, generating a 555 bp amplicon (Rec2 RT-PCR; Supplementary table 1). Primers were subsequently modified by the 5’ addition of Illumina pre-adapter and index primer binding sequences to enable high-throughput Illumina sequencing (Rec2.tailed RT-PCR; Supplementary table 1). Assays were experimentally validated to confirm reactivity with all known Australian lagoviruses. For a subset of samples (prior to the development of the Rec2 RT-PCR), amplification over the RdRp-VP60 junction was performed with alternative primer sets (Supplementary table 1). Briefly, each 25 µl PCR reaction comprised 1x OneStep Ahead RT-PCR master mix (Qiagen), 1x OneStep Ahead RT mix (Qiagen), 0.5 µM each primer, and 1 µl of extracted RNA diluted 1/10 in nuclease-free water. Cycling conditions were: 45 °C for 15 min, 95 °C for 5 min, followed by 40 cycles of 95 °C for 15 sec, 55 °C for 15s, 68 °C for 2 min, with a final extension of 68 °C for 5 min.

**Table 1.**
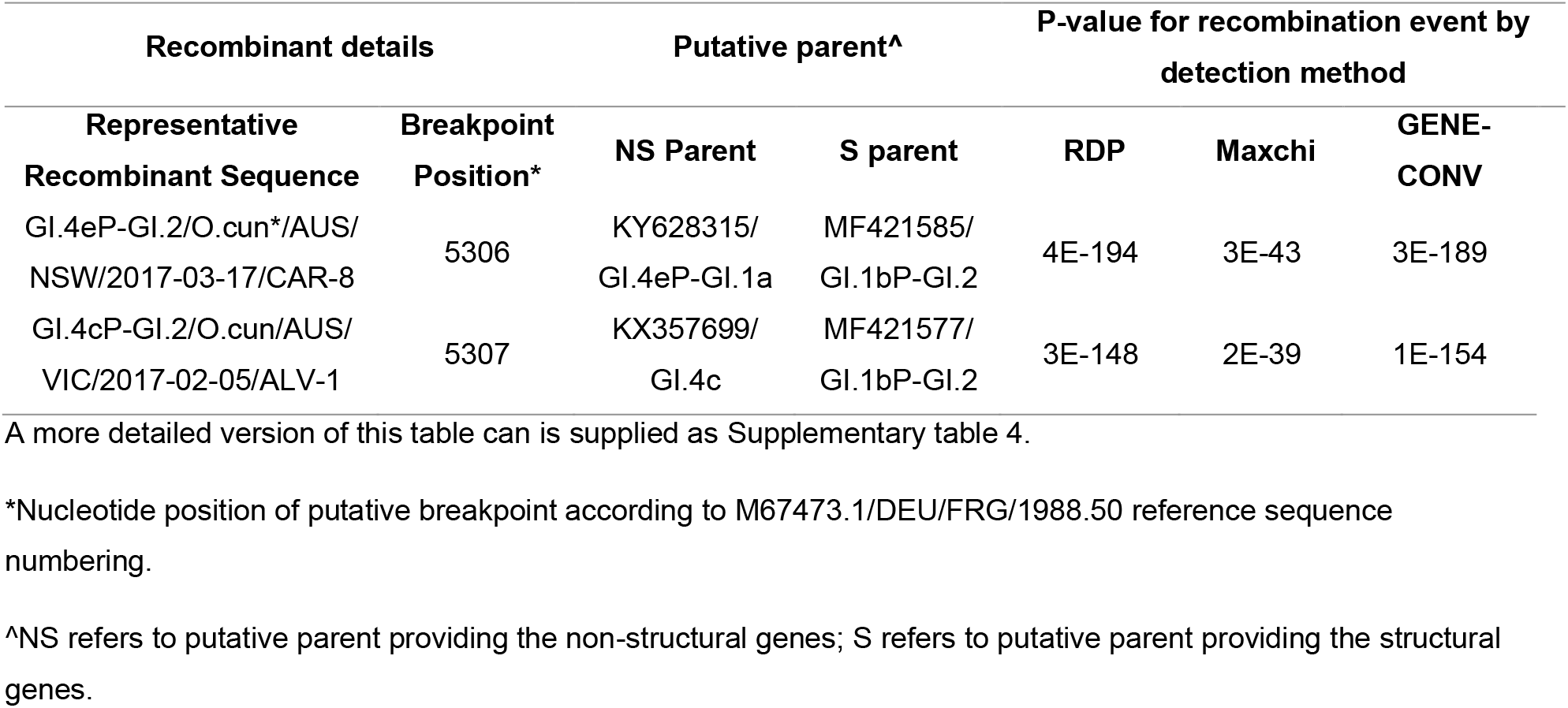
Recombination analyses.

Amplicons were purified and either sent for Sanger sequencing at the Biomolecular Resource Facility, The John Curtin School of Medical Research, Australian National University, or were indexed, pooled, and sequenced on an Illumina MiSeq (300-cycle v2 chemistry) according to the manufacturer’s directions. After quality trimming, sequences were partitioned into regions upstream and downstream of the RdRp-VP60 junction. Each partition was aligned with representative sequences of known Australian lagovirus variants using MAFFTv7.450 [41] in Geneious Prime 2020.0.4, and an RdRp and VP60 variant type was assigned for each sample. Four isolates were identified as mixed infections based on sequencing. Alignments of partial sequences are available at https://doi.org/10.25919/758f-4t15.

### Molecular epidemiological analyses

To investigate the geographical distribution and relative incidences of recombinant variants over time, positive samples were restricted to those reported after 1 January 2016; i.e., 6 months prior to the first detection of a novel recombinant (n = 1,139). Samples identified as RHDVa-K5 (n = 121) were excluded from further analyses since these were mostly associated with release sites of this approved biocontrol virus; that is, this variant was not transmitting through rabbit populations and was therefore not competing epidemiologically with other virus variants [39]. Samples from the ACT were grouped with those from NSW, since the ACT is a small jurisdiction (2,358 km^2^) located within NSW and was considered epidemiologically to be part of NSW.

For the 4c-recombinant lineage designation, 169 4c-recombinant sequences spanning the RdRp-VP60 junction were aligned with 77 4c-recombinant full genomes and 26 publicly available GI.4 genomes using MAFFTv7.450 [41]. The taxa included in the final data set are detailed in Supplementary table 2. The alignment was trimmed to nucleotide positions 5,115 – 5,619, based on the M67473.1 DEU/FRG/1988.50 reference sequence (https://www.ncbi.nlm.nih.gov/genbank/). The remaining 109 4c-recombinant sequences obtained in this study were not suitable for inclusion in the final alignment because they were generated either using Illumina sequencing or an alternative primer set and thus did not span the complete region of the alignment. Model selection and maximum likelihood (ML) tree inference was conducted using IQ-TREEv1.6.11 [42], and branch supported was estimated by 1,000 replicates of the SH-like approximate likelihood ratio test [43]. The tree was rooted on the internal branch leading to the RCV-A1 lineage.

### Full genome sequencing

Of the 1,034 samples genotyped to the variant level, a subset of 224 were selected for full genome sequencing (Supplementary table 3). These samples were selected to be geographically and temporally representative, with a focus on the novel 4e- and 4c-recombinants. Viral genomes were amplified in overlapping fragments, and DNA libraries were prepared and sequenced using Illumina Miseq technology as described previously [35, 40]. Primers used for amplification of the overlapping fragments are detailed in Supplementary table 1. Consensus sequences were constructed by mapping cleaned reads to the lagovirus GI reference sequence (GenBank accession M67473.1 DEU/FRG/1988.50) using Geneious Prime 2020.0.4. Sequences were deposited in GenBank under accession numbers MW460205 – MW460242.

### Recombination analysis

To further characterise the putative novel 4c-recombinant variant, and the newly sequenced GI.4e-recombinants, recombination analyses were conducted on full genome sequences using the RDP4 software [44]. The complete genome alignment included lagovirus GI potential parent sequences from GenBank (https://www.ncbi.nlm.nih.gov/genbank) (n = 384, 7,309 nt, Supplementary table 4). Recombination was reported if detected by at least two of three primary scanning methods (RDP, MaxChi and GENECONV), with a highest acceptable p-value of 0.05 with Bonferroni multiple comparison correction. The BootScan, 3Seq, CHIMAERA, and SciScan methods were used to verify recombination events identified using the primary methods.

### Phylogenetic analyses

Newly sequenced lagovirus genomes were aligned with representative GI lagovirus sequences from GenBank (https://www.ncbi.nlm.nih.gov/genbank) using MAFFTv7.271 [41]. The complete genome alignment was split into two data sets representing (i) the NS genes (n = 240 sequences, 5,231 nt, Supplementary table 5), and (ii) the VP60 structural gene (n = 272, 1,740 nt, Supplementary table 6). An additional VP60 alignment was constructed that contained the VP60 of the newly sequenced viruses along with all published Australian RHDV2 VP60 sequences (n = 332, 1,737 nt, Supplementary table 7). Model selection and ML tree estimation was performed using IQ-TREEv1.6.11 [42], as described above. Ancestral state reconstruction to identify nonsynonymous changes along individual branches was carried out using PAMLv4.9 [45].

### Phylogeographic analyses

A Bayesian Markov chain Monte Carlo (MCMC) approach was employed to infer time-scaled phylogenies, and from this, to infer the temporal pattern and most likely geographic location of internal nodes (i.e., phylogeography). All data sets were initially screened using TempEst [46] to ensure that sufficient temporal signal was present, using ML phylogenies as input to construct linear regressions of root-to-tip genetic distances against sampling time. Due to recombination between the NS and capsid genes, two data sets either side of the recombination breakpoint were analysed separately: (i) a VP60 gene data set containing all published Australian GI.2 capsid sequences together with the capsid genes of the recombinants sequenced here (n = 332, 1,737 nt, Supplementary table 7); and (ii) an NS genes data set of all published Australian GI.4 sequences together with the NS genes of the recombinants sequenced here (n = 188, 5,218 nt, Supplementary table 8). The Bayesian Evolutionary Analysis by Sampling Trees (BEAST) software v1.8 [47] was used to conduct Bayesian MCMC analysis of each data set, using substitution models inferred using ModelFinder as implemented in IQ-TREEv1.6.11 [42] (VP60, SYM+G4; NS, SYM+I+G4) [48]. A discrete traits partition indicating the sampling location (Australian state) was included to facilitate ancestral state reconstruction of location (utilizing the symmetric substitution model, inferring social network with Bayesian Stochastic Search Variable Selection, and a strict clock model). All analyses were run twice to convergence (defined as an effective sample size >200) to confirm consistency. Marginal likelihood estimation using path sampling/stepping-stone sampling was used to assess the most appropriate clock model prior (strict vs uncorrelated log-normally distributed [UCLD]) and tree prior (Gaussian Markov random field Bayesian skyride model vs constant size coalescent vs exponential coalescent) for the nucleotide partition.

The NS data set had a better temporal signal (R-squared of 0.86 on linear regression of root-to-tip genetic distance against sampling time) and a larger sampling window compared to the VP60 data set and was therefore considered the most informative. The UCLD clock model and constant population size coalescent tree prior had the highest marginal likelihood for this data set.

### Figures

Figures were constructed using R v4.0.3 [49] using the following packages: ‘ggtree v2.2.1’ [50], ‘treeio v1.12.0’ [51], ‘scales v1.1.1’ [52], ‘tidyverse v1.3.0’ [53], ‘ape v5.4’ [54], ‘phytools v0.7-47’ [55], ‘phangorn v2.5.5’ [56], ‘cowplot v1.0.0.9000’ [57], ‘lubridate v1.7.8’ [58], ‘plyr v1.8.6’ [59], and ‘ozmaps v0.3.6’ [60]. Figure 3B was generated in GeneiousPrime 2020.2.4 using 3-D structure viewer.

**Figure 3:**
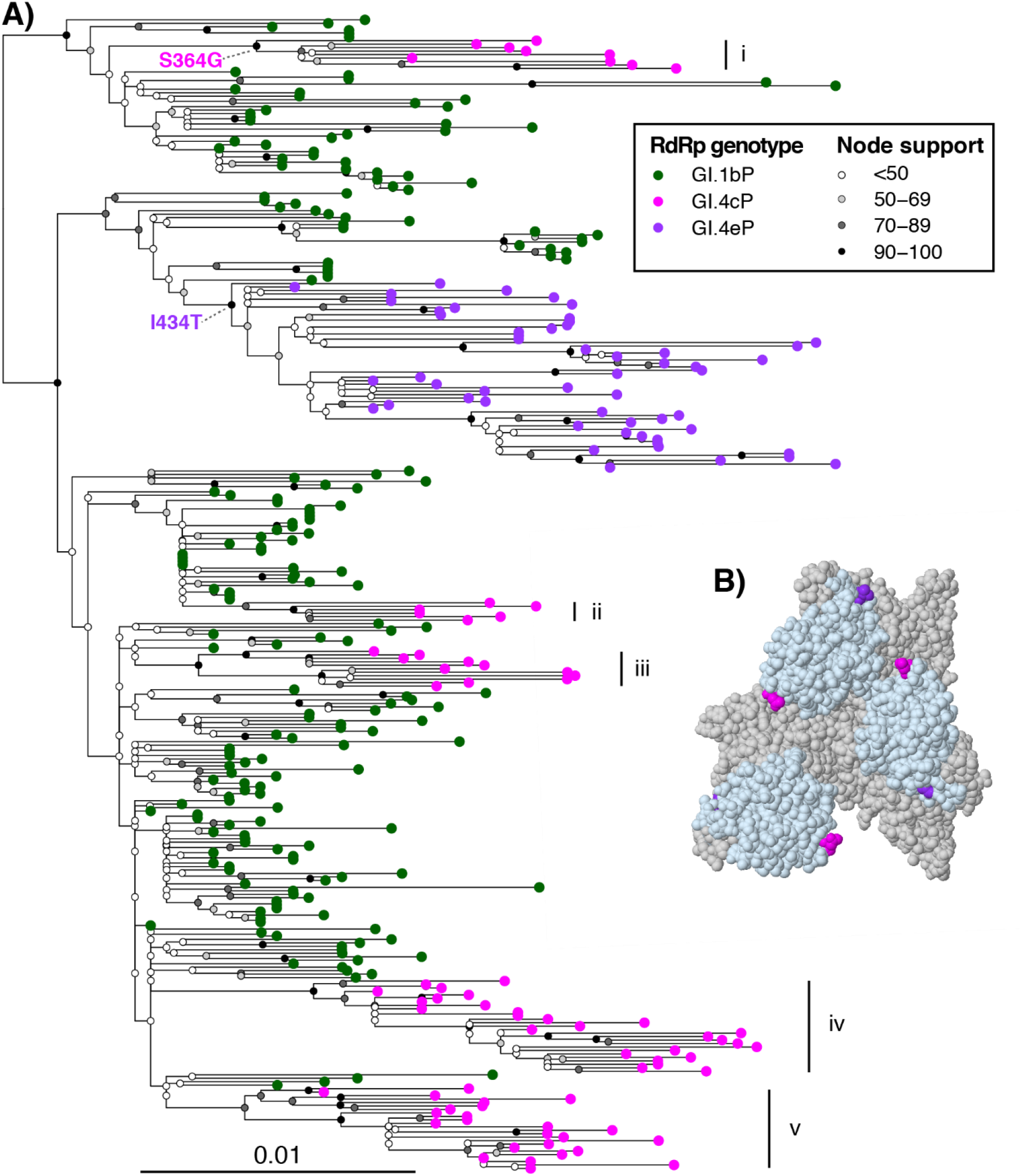
ML phylogenetic tree of Australian GI.2 VP60 sequences and amino acid changes associated with the emergence of 4c- and 4e-recombinant variants. (A) A phylogeny was inferred from the VP60 gene sequences (n = 332) of newly sequenced Australian GI.2 lagoviruses (including recombinants) along with representative published sequences. Coloured tip points represent the polymerase (RdRp) genotype: GI.1bP, green; GI.4cP, pink; GI.4eP, purple. GI.4cP lineages are labelled (i through v). The phylogeny was midpoint rooted and the scale bar represents nucleotide substitutions per site. Non-parametric bootstrap support values (1,000 replicates) are indicated by shaded circles at the nodes. Associated metadata including taxon names are provided in Supplementary table 7. Ancestral state reconstruction to identify nonsynonymous changes associated with internal divergence events was conducted using PAML. All changes occurring on branches leading to the GI.4cP and GI.4eP lineages are labelled at the relevant node. (B) These changes were mapped to an asymmetric unit of the atomic model of lagovirus GI VP60 (PDB accession: 3J1P). The P2 domain is coloured in light blue. Amino acid 364, associated with the change in 4c-recombinant lineage i, is coloured pink and 434, associated with the 4e-recombinant, is coloured purple.

## Results

### Phylogenetic analyses reveal five independent recombination events in GI.4cP-GI.2 viruses

A recombinant lagovirus variant (GI.4eP-GI.2; 4e-recombinant) was previously detected in Australia using a multiplex RT-PCR assay. This assay is only able to detect specific recombinant variants [31]. Therefore, to comprehensively screen for lagovirus recombination events we sequenced the RdRp-VP60 junction region (i.e., the typical recombination breakpoint) of 1,034 lagoviruses collected between January 2014 and September 2020 (inclusive) from wild and domestic lagomorphs found dead across Australia. This screen identified six distinct lagovirus variants circulating in Australia during the study period: four known variants (GI.1cP-GI.1c (RHDV1), GI.1bP-GI.2 (RHDV2), GI.1aP-GI.1a (RHDVa-K5), GI.4eP-GI.1a (RHDVa-Aus)), the previously reported 4e-recombinant (GI.4eP-GI.2), and a novel GI.4cP-GI.2 putative recombinant (4c-recombinant).

We selected a subset of novel 4c-recombinant viruses, recently emerged 4e-recombinant viruses, and parental RHDV2 viruses for full genome sequencing, selecting isolates that were representative of the temporal and geographical spread of these variants. Recombination analyses of newly sequenced 4e-recombinant viruses confirmed these as recombinants with a breakpoint at the RdRp-VP60 junction, supporting our previous finding (Table 1, Supplementary table 4). Recombination analyses of the novel putative 4c-recombinant viruses also detected a recombination breakpoint at the typical RdRp-VP60 junction (Table 1). The most likely parental variants for the 4c-recombinant, as determined by the RDP4 software, were benign Australian GI.4 (RCV-A1) lagoviruses in the NS region (GenBank accession KX357699) and Australian GI.2 (RHDV2) viruses in the VP60 region (GenBank accession MF421577), strongly suggesting that this recombination event occurred within Australia after the arrival of GI.2 in late 2014.

Subsequently, the nucleotide alignment was partitioned either side of the putative breakpoint and ML phylogenetic trees were inferred for the NS genes and the VP60 gene separately. There was clear incongruence between the NS and VP60 phylogenies, supporting the results of the RDP4 analysis (Figure 2). Strikingly, ML phylogenies based on the VP60 region revealed five distinct lineages of the 4c-recombinant variant, suggesting multiple independent emergences of this variant (Figure 3A). In contrast, the NS gene sequences from this variant formed a monophyletic group (collapsed in Figure 2A). All 4e-recombinant sequences fell within a single clade in both trees (Figure 2).

**Figure 2.**
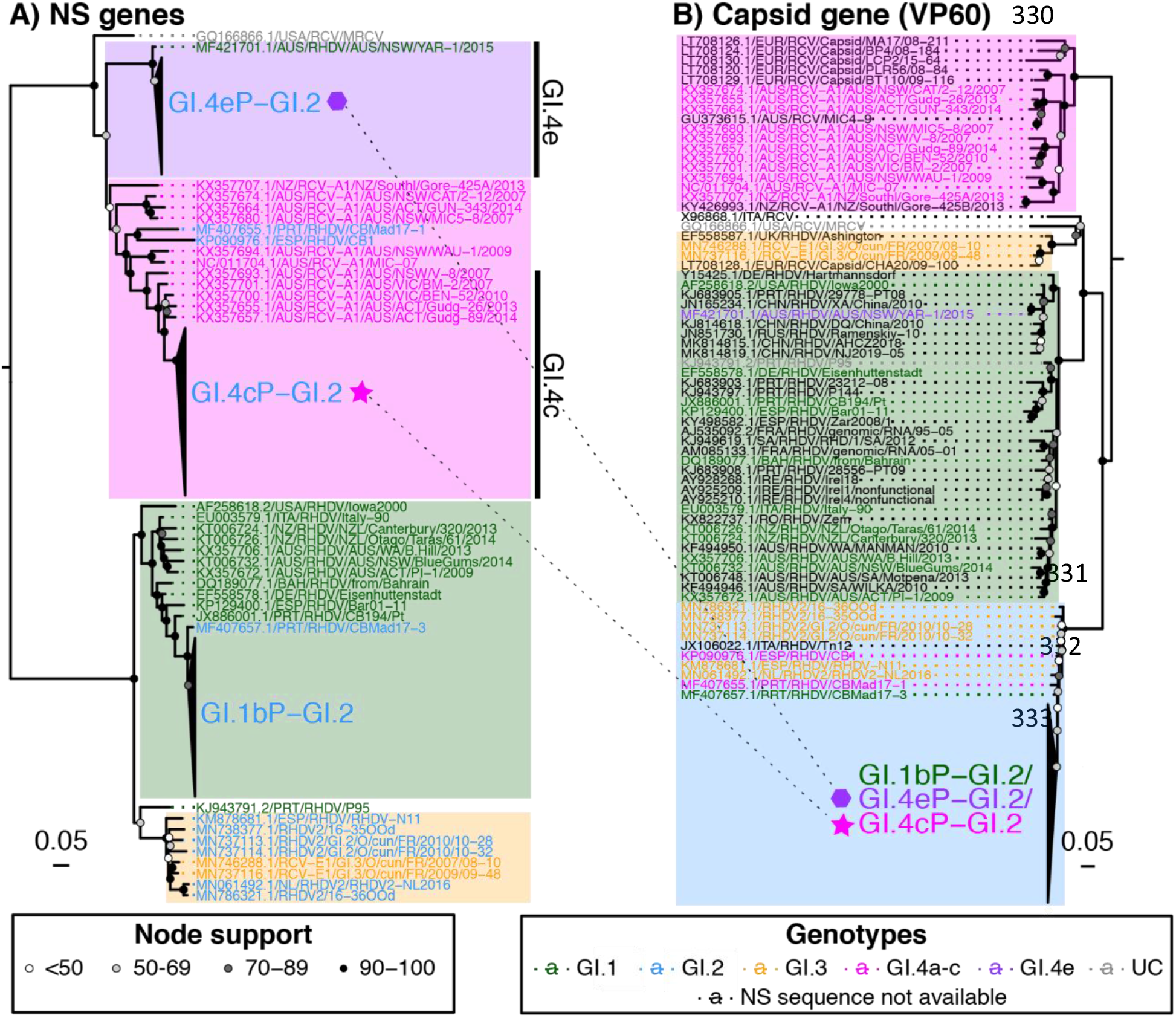
ML phylogenetic trees of the (A) NS genes and (B) the VP60 gene of representative lagovirus sequences and the Australian recombinant sequences. Discrepancies between the highlight and text colour indicate recombinant sequences. Clades are highlighted according to the genotype of the genetic segment analysed in the tree, while taxa names are coloured by the genotype of the alternative segment (i.e., taxa names in the NS gene tree are coloured by the genotype of the capsid gene, and vice versa). Taxa names coloured black indicate that the NS sequence data is unavailable. UC, unclassified. Major clades have been collapsed for visualization purposes. All Australian recombinant sequences from this study fall within the collapsed clades indicated by the star (GI.4cP-GI.2) or hexagon (GI.4eP-GI.2) symbols. The dotted lines between the two trees link the Australian recombinant clades. Phylogenies were midpoint rooted and the scale bars represent the number of substitutions per site. Bootstrap support values (1,000 replicates) are indicated by shaded circles at the nodes.

### Rapid epidemiological replacement of parental RHDV2 by the newly emerged 4c- and 4e-recombinants

Detections of the new 4c-recombinant variant consistently outnumbered those of the parental RHDV2 in VIC by April 2018, 14 months after the first detection in Alvie, VIC, on 5 February 2017, suggesting that this variant was outcompeting previously circulating viruses (Figure 4). This pattern of emergence and replacement was also observed in TAS (Figure 4). This variant was detected sporadically in NSW/ACT from October 2017 where it cocirculated with the locally dominant 4e-recombinant; notably, the proportional incidence of the 4c-recombinant variant has increased in this region during 2020. Interestingly, a single detection of the 4c-recombinant was identified in WA in November 2018 in two domestic rabbits recently imported from TAS as breeding stock. The sequence of this 4c-recombinant virus detected in WA clearly nested within the genetic diversity found in TAS (see Figure 4A, clade v), suggesting a single incursion into WA with no ongoing local transmission. During the study period, the novel 4c-recombinant was recovered from several juvenile animals less than 800 g bodyweight and as young as 4 weeks old, demonstrating pathogenicity in young rabbits, like all other GI.2 variants characterized so far. It was also identified in one European brown hare (*Lepus europaeus*).

**Figure 4:**
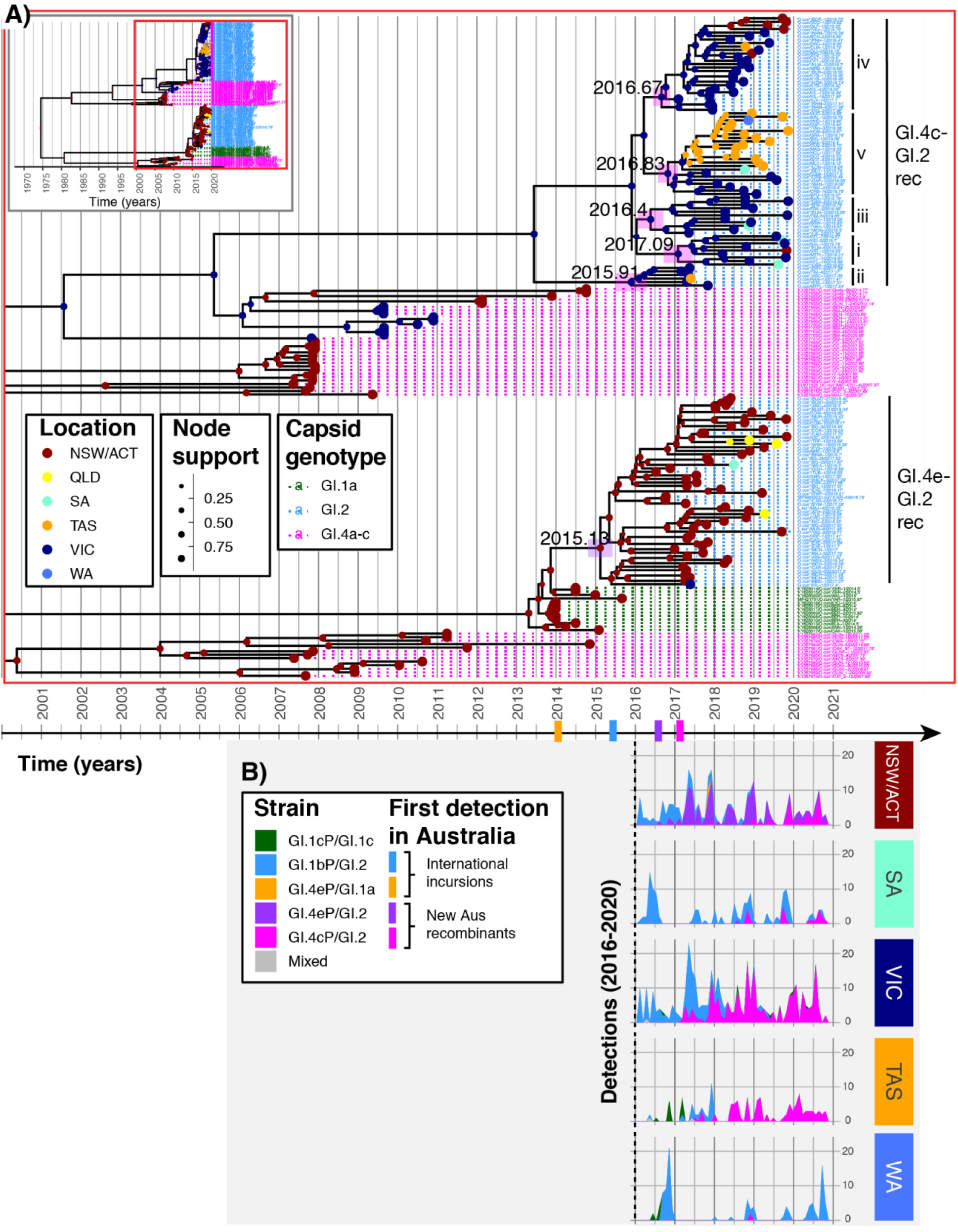
Timing and location of the emergence of Australian lagovirus recombinants based on time-structured phylogeographic reconstruction and RT-qPCR detections. Panels A and B are scaled to the same x-axis (shown between the two panels), given as time in years. (A) Time-structured phylogeographic analysis of Australian GI.4 lagovirus NS genes (uncorrelated log-normally distributed clock model, constant tree prior). The inset indicates the section of the phylogeny shown. Node and tip points are coloured according to location (Australian state) and node points are sized according to posterior support for that clade. Taxon names are coloured by capsid variant. The mean time to most recent common ancestor for the GI.4eP-GI.2 and each GI.4cP-GI.2 recombinant lineage is indicated at the respective internal node, and horizontal bars at these nodes represent the 95% highest posterior density (HPD). The recombinant (rec) clades are labelled, including the GI.4cP-GI.2 lineages (i – v). (B) Lagovirus positive samples collected in NSW/ACT, SA, VIC, TAS, and WA from 2016 to 2020 (n = 739) were genotyped to the variant level by sequencing either side of the typical calicivirus recombination breakpoint. The number of detections of each variant by month are shown for each geographical region as an area plot, with the plotted area coloured by variant.

We also examined the distribution of the previously reported 4e-recombinant variant [31]. Most detections (n = 140) were identified in NSW/ACT. The earliest detection was in Tubbul, NSW, on 13 July 2016, and by March 2017 it had mostly replaced the parental RHDV2 to become the dominant lagovirus variant in this region (Figure 4B). More recently, in late 2019 and throughout 2020, it has cocirculated with the 4c-recombinant in NSW/ACT. Although most detections were in adult rabbits, the 4e-recombinant was also recovered from three European brown hares and from several juvenile rabbits down to 250 g bodyweight, suggesting a similar host tropism to the parental RHDV2 virus and the novel 4c-recombinant.

### All recombination events occurred within a two-year timeframe in eastern Australia

To determine the temporal dynamics, the location of emergence, and the rate of viable recombination events in Australian lagoviruses, we conducted time-structured phylogenetic analyses for the NS and capsid genes of recombinant viruses and related sequences (i.e., Australian GI.2 VP60 and GI.4 NS sequences). Based on regression analysis, data from the GI.4 NS dataset was considered more accurate and is therefore presented in Figure 4. Based on these analyses, the 4e-recombinant was the first variant to emerge, with the most recent common ancestor of this lineage dated to early to mid-2015 (95% highest posterior density [HPD] NS dataset, 2014.8 – 2015.43) (Figure 4A). For the 4c-recombinants, the 95% HPD intervals of the time to most recent common ancestor (TMRCA) overlapped between lineages, making it difficult to confidently define the exact timings and order of emergence of each lineage (Figure 4A). However, the data clearly show that these five recombination events all occurred within the space of two years. This two-year timeframe was consistently observed regardless of whether the analysis was conducted on the NS genes GI.4 data set or the VP60 gene GI.2 data set. Our phylogeographic analysis (Figure 4A) suggests that all the 4c-recombinant lineages emerged in VIC (probability >0.99 for all lineages), with subsequent spread to other states. Notably, this was observed even for lineage v, which was first detected in Tasmania. In contrast, the 4e-recombinant clearly emerged in NSW/ACT (probability >0.99, Figure 4A); this is further supported by our epidemiological data, where we found only limited detections of this variant in other states (QLD, VIC, SA).

### Continued cocirculation of 4c-recombinant lineages

The 4c-recombinant viruses were initially assigned to lineages based on the GI.2 VP60 phylogeny (Figure 3A). For GI.4c-recombinant viruses where a full genome sequence was not available, we first inferred a phylogeny from a 504 nt region spanning the RdRp-VP60 junction (Figure 5); this phylogeny was annotated using taxa for which the lineage was definitively known from the VP60 phylogeny. While sequences from each 4c lineage (as defined from the complete genome sequences) did not form clades with individual common ancestors in the RdRp-VP60 junction phylogeny, they did form distinct groups with visible genetic distance between them (Figure 5). These groups were distinct enough to assign a 4c lineage type to those samples with partial sequence only. After assigning lineages to all GI.4c-recombinants, we explored the interactions between the different lineages by mapping their sampling locations over time. We found that four of the five lineages were already circulating in 2017 (Figure 5). Lineage ii was not detected after 2017 while the other four lineages continued to be detected throughout 2018, 2019, and 2020. In VIC, where all lineages are circulating, there was no clear indication of geographical clustering of the different lineages. In TAS, lineages ii, iii, and iv were detected sporadically while lineage v has remained dominant over time, suggestive of a founder effect in this region. Similarly, in NSW/ACT lineage iv has remained dominant, also suggesting a regional founder effect, while lineages i, ii, and iii were only detected intermittently.

**Figure 5:**
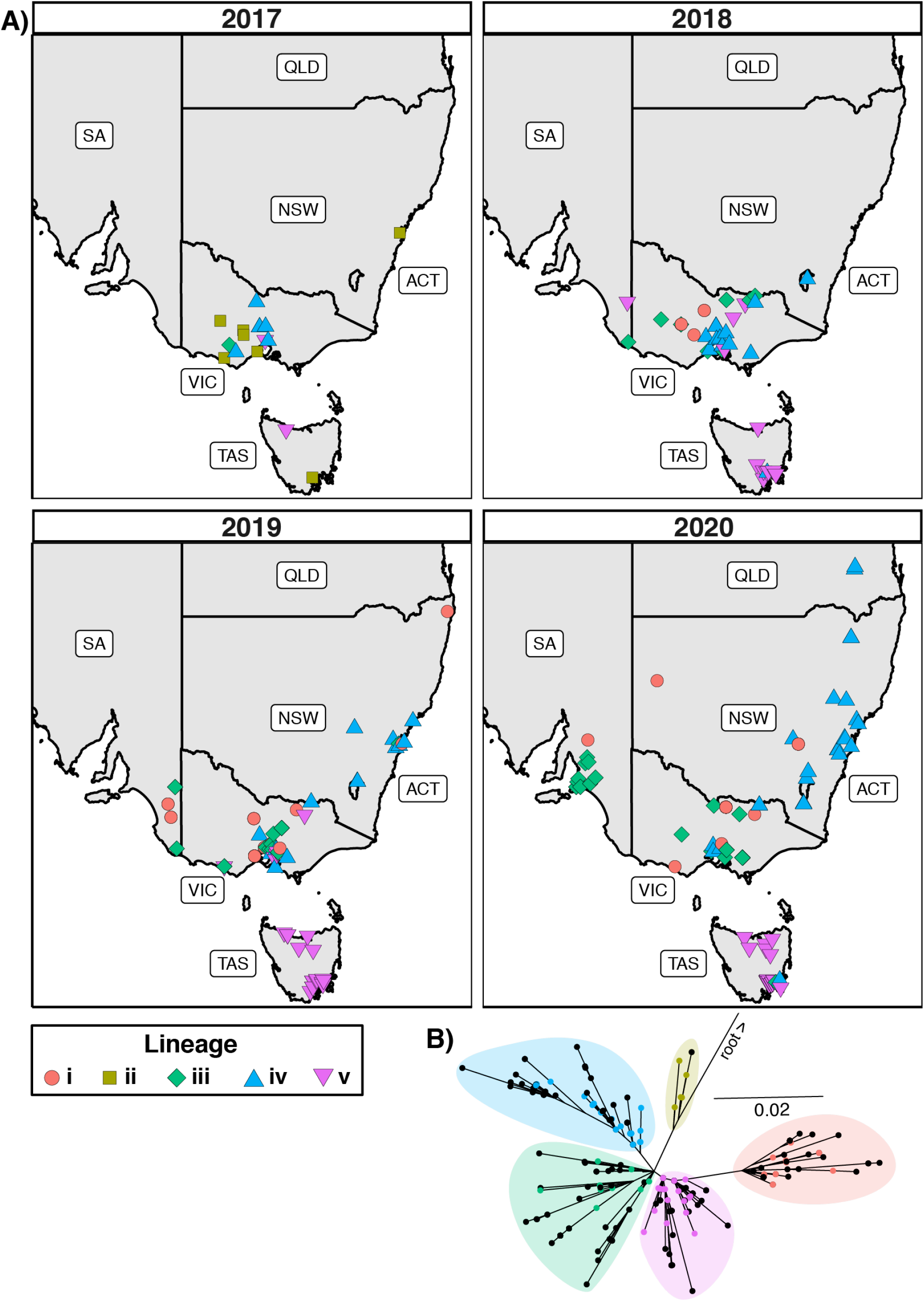
Geographical distribution of 4c-recombinant lagoviruses in Australia, 2017 to 2020. (A) Sampling locations for the 4c lineages were mapped for each year following the emergence of this variant in 2017. (B) 4c-recombinant viruses were allocated to a 4c lineage based on ML phylogenetic grouping of the partial RdRp-VP60 sequence (504 nt) with reference sequences. Reference sequences were annotated based on full genome sequencing and are indicated in the tree by the coloured tip points. The phylogeny was midpoint rooted and the scale bar represents nucleotide substitutions per site.

### The emergence of 4c-recombinant variants was not associated with antigenic changes in the VP60 capsid protein

To determine whether the observed epidemiological replacement of 1) the parental RHDV2 virus by the novel 4c-recombinant in VIC and TAS, and 2) the 4e-recombinant in NSW/ACT were associated with nonsynonymous, potentially antigenic, changes in the VP60 capsid protein we performed ancestral state reconstruction using our prior Australian GI.2 VP60 phylogeny. Notably, four of the five 4c-recombinant lineages showed no amino acid changes in the capsid protein relative to the inferred ancestral RHDV2 node, suggesting that the emergence of these lineages was not associated with changes in antigenicity (Figure 3A). However, nonsynonymous mutations were identified in 4c-recombinant lineage i and in 4e-recombinant viruses (Figure 3A). When we mapped these mutations back to an atomic model of lagovirus GI VP60, both mutations were within the protruding domain, specifically the P2 subdomain, which is known to contain immunodominant epitopes (Figure 3B) [61]. The 4c-recombinant lineage i was associated with an S364G mutation, while an I434T mutation was present in all lineage 4e-recombinants (Figure 3A). Thus, it is possible that the epidemiological fitness of the 4e-recombinant variant in NSW/ACT may be attributable to antigenic changes in the capsid; however, this cannot explain the replacement of parental RHDV2 by 4c-recombinant variants in VIC or TAS, or the continued co-circulation of 4c-recombinant lineages without changes in VP60. Furthermore, in NSW/ACT the 4e-recombinant variant, despite the I434T change in VP60, appears to be currently undergoing replacement by the 4c-recombinant.

## Discussion

In a global landscape where population immunity to GI.1 and GI.1a viruses was widespread in lagomorphs, it is unsurprising that GI.2 viruses were so successful and rapidly replaced GI.1 strains. We were interested to determine whether, after establishment, GI.2 viruses would continue to evolve via immune selection and antigenic drift or by other mechanisms. We were also interested to document whether variants without the GI.2 capsid would emerge as population immunity increased against this genotype. Instead, we found that the GI.2 capsid was retained and that the emerging recombinant variants acquired alternative NS genes. It is becoming increasingly clear that recombination is a major driver of calicivirus diversity, facilitating the emergence of new variants including those with pandemic and panzootic potential [24, 25, 62, 63]. However, the frequency at which viable recombination occurs and the genetic drivers of epidemiological fitness are still poorly understood. In this study, a continent-scale natural competition experiment of lagoviruses in Australia, we found that viable recombination occurs at an extremely high frequency given the right agent, host, and environmental circumstances.

### Six viable recombination events in Australian lagoviruses between 2014 and 2018

Shortly after detecting two exotic lagovirus incursions in Australia, RHDVa-Aus (GI.4eP- GI.1a) in 2014 and RHDV2 (GI.1bP-GI.2) in 2015 [32, 37], we reported the emergence of a recombinant variant of these two viruses (4e-recombinant; GI.4eP-GI.2), detected in July 2016 in NSW [31]. This finding prompted the development of improved molecular tests for the detection of recombination and the subsequent retrospective and prospective screening of lagovirus-positive samples for further recombination events.

This screen identified an additional novel recombinant lagovirus, a GI.4cP-GI.2 variant (4c-recombinant), first detected in VIC in February 2017. Further analysis determined that the VP60 genes of this 4c-recombinant separated into five distinct clades (Figure 3A), suggesting that this variant emerged on five separate occasions following at least five independent recombination events. Recombination was already known to occur in lagoviruses. For example, in the last decade, the lagovirus GI.2 capsid has been detected in combination with five different sets of NS genes [24, 27, 31, 33]. Our study demonstrates that viable recombination during natural infection occurs even more frequently than previously appreciated but may often go undetected if different events result in closely related progeny. Since the first detection of RHDV2 in Australia in mid-2015 [37] this capsid gene has recombined and produced epidemiologically viable variants at least six times in a two-year period.

The previously reported 4e-recombinant arose from a recombination event between RHDVa-Aus (GI.1aP-GI.2) and RHDV2 (GI.1bP-GI.2), both recombinant viruses themselves that were exotic incursions detected in Australia in January 2014 and May 2015, respectively, although the source of these viruses remains unknown [31, 32, 37]. Detections of the parental RHDVa-Aus were predominantly restricted to the Sydney basin region of NSW, with two cases detected in QLD and one further case in a wild rabbit in the ACT [32]. Therefore, it is not surprising that the 4e-recombinant emerged in NSW/ACT, according to our epidemiological data and phylogeographic analysis (Figure 4). The 4c-recombinants arose from recombination events between RHDV2 (GI.1bP-GI.1) and endemic RCV-A1 (GI.4c) viruses [14, 37]. Emergence of the 4c-recombinant variants in the south-eastern states of Australia is consistent with high rabbit population densities and a high seroprevalence to GI.4 viruses in these regions compared to more arid regions of Australia [64].

Despite the VP60 phylogeny suggesting multiple recombination events between GI.4c and GI.2 viruses, only a single unique recombination event was detected by the RDP4 program. This is not surprising given that the breakpoint locations are indistinguishable and these recombinants (and their true parents) are very closely related. Indeed, it was only possible to infer the true number of recombination events because of the high sampling frequency of parental RHDV2 viruses, which has been maintained for this and previous studies. Had RHDV2 sequences from intervening clades not been sampled, the 4c-recombinants would have formed a monophyletic clade and a single recombination event assumed. This is demonstrated by the NS genes phylogeny (Figure 4A), in which the 4c-recombinant sequences form a single clade within the greater GI.4c clade. Importantly, within this clade the sequences group into the same five lineages as seen in the VP60 tree. Compared to pathogenic GI.2 viruses, benign RCV-A1 (GI.4c) viruses are greatly under-sampled in Australia, with the most recent RCV-A1 sequences (n = 6) derived from samples collected from the ACT in 2012 and 2014 [35]. Aside from three sequences sampled in 2014, no other RCV-A1 sequences were available for the entirety of the study period. Consequently, a close relative of the RCV-A1 parental virus has not been sequenced, suggesting a hidden diversity of unsampled RCV-A1 viruses in Australian rabbits. This study provides a lower bound to the rate of recombination in these viruses; it is possible that additional recombination events have occurred with GI.4c viruses that are not observable against the current background of available GI.2 VP60 sequences.

### Lagoviruses show frequent recombination between the GI.2 capsid and GI.4 NS sequences

Interestingly, all the recombination events detected during the study period involved the pairing of a GI.2 capsid and the NS genes of a GI.4 virus. Despite screening over 800 lagovirus-positive samples, we found no evidence for recombinant variants with either capsid or NS sequences of GI.1 viruses.

For recombination to occur several criteria must be met [23]. Firstly, there must be coinfection of an individual host. This is influenced by host tropism, the prevalence of each parental virus in the population, and the duration of infection. Secondly, there must be coinfection of a single cell, both viruses must replicate within this cell, and precise template switching must occur to generate viable gRNA. Finally, the resultant variant must be epidemiologically competitive. That is, it must be able to successfully transmit, to establish infection in new hosts, and to avoid being outcompeted by either the parental or other circulating variants.

Following the epizootic incursion of an antigenically novel virus, RHDV2, into a naïve population, the prevalence of this variant was extremely high in Australian rabbits, reflected by the estimated population-wide mortality rate of 60% following the arrival of RHDV2 [39]. The detection of four GI.1 and GI.2 co-infections in this study between 2015 and 2017 demonstrates that both viruses were circulating at sufficiently high prevalence for mixed infections to occur. However, this doesn’t explain the predilection for GI.4 (RCV) NS sequences. The discovery of 4c-recombinants was surprising, since the parental RCV-A1 is a benign enterotropic lagovirus [14], while RHDV2 (and RHDV1) viruses are virulent and hepatotropic. This would seem to preclude coinfection of individual cells with both viruses. Yet, several recombinants of RHDV2 and benign enterotropic RCVs have also been reported from Europe [64]. This suggests that active replication of both RCVs and RHDV2 must be occurring in the same target cell. Extrapolating from our current understanding of human and murine norovirus tropism, macrophages or other immune cells may be likely candidates [65]. This is further supported by the detection of plus- and minus-strand viral RNAs in splenic and alveolar macrophages of rabbits experimentally infected with RHDV1 [66]. Additionally, RCVs typically infect young rabbits early in life [12, 14]. Although young rabbits can be infected with RHDV1 viruses, robust innate immunity limits the extent of viral replication in an age-dependent manner [34, 39]. Thus, mixed infections with RHDV1 and RCVs are probably infrequent. This age-dependent resistance is not observed with RHDV2 infection [67]. Furthermore, the duration of infection is longer for benign RCVs compared to pathogenic variants, where infected individuals typically die within 48 – 72 hours post-infection [2]. These factors may at least partly explain why RCV-RHDV2 recombinants appear to emerge more frequently than RHDV1-RHDV2 recombinants.

### Epidemiological drivers of lagovirus emergence and spread

Both the 4e-recombinant and 4c-recombinant rapidly replaced the dominant circulating parental RHDV2 in NSW/ACT and VIC/TAS, respectively. This replacement, at least for several 4c-recombinant lineages, occurred without any associated antigenic changes in the capsid protein, demonstrating that the replacement was not driven by antigenic escape. This shows that NS sequence variation is an important driver of epidemiological fitness in lagoviruses, complementing similar findings in human and murine noroviruses [68, 69]. For example, the pandemic GII.P16/GII.4 Sydney 2012 norovirus, which does not contain unique substitutions in the capsid, has substitutions within the RdRp that are proposed to increase transmissibility [69]. Infection with this variant also resulted in increased viral shedding compared to other norovirus genotypes, as measured by higher faecal viral loads [70]. Previous studies have shown that RdRp fidelity and intra-host viral diversity affect transmissibility of murine norovirus *in vivo*, with high-fidelity variants being less efficiently transmitted than the wild-type variant [71]. Taken together, these studies demonstrate that NS proteins, particularly the RdRp, are important drivers of calicivirus fitness. Interestingly, the *in vitro* polymerase replication activity of the cloned GI.4c RdRp was previously shown to be at least two times that of the RHDV1 (GI.1c) RdRp [72], although comparison to the GI.1b RdRp, the specific ‘competing’ variant in the current study, was not reported. Analogous to the findings in murine norovirus, we propose that a higher replication rate of GI.4 RdRps may lead to increased intra-host viral diversity and transmissibility of GI.4 recombinants. However, we cannot rule out that other NS proteins may also contribute to the observed high fitness of the recombinant variants.

Within the 4c-recombinants, there was no evidence of dominance of any one lineage over time. The apparent dominance of lineage iv in NSW/ACT and lineage v in TAS is consistent with founder effects in both of those states. This demonstrates that different lagoviruses can cocirculate at relative equilibrium over extended periods of time. This further supports that the observed rapid replacement of parental RHDV2 (GI.1bP-GI.2) by GI.4P recombinant variants in this study is due to a fitness advantage of these variants, conferred by the NS proteins and possibly associated with the RdRp.

### The lagovirus capsid governs host and tissue tropism and is correlated with virulence

Both the newly identified 4c-recombinant and the previously emerged 4e-recombinant are virulent, hepatotropic viruses that were recovered from the livers of both rabbits and hares and from rabbits of all ages in this study. This tropism mimics that seen with other GI.2 viruses [73–80]. In stark contrast, the parental RCV-A1 is a benign, enterotropic virus that has only been recovered from rabbits [14], while the RHDVa-Aus variant, although virulent and hepatotropic, has only been found in adult rabbits. Our findings suggest that it is the lagovirus capsid that confers both host and tissue tropism and that tissue tropism is correlated with virulence.

With the broader host tropism conferred by the GI.2 capsid, there is increased potential for the emergence of novel epizootic lagovirus variants through both intragenotypic and intergenotypic recombination. Hares are known to carry their own, presumed benign, caliciviruses [16–18, 31, 33, 73–79]. Indeed, the first intergenotypic lagovirus recombinants were recently reported from Germany [33]. Since the incursion of RHDV2 into North America in 2020 this variant has also been reported to infect several *Sylvilagus* species (cottontail rabbits) [81]. Although endemic *Sylvilagus* calciviruses have never been reported, very limited sampling has been conducted in this species. It remains to be seen whether North American leporids may be a new reservoir for the emergence of novel lagoviruses with panzootic potential. This highlights the need for ongoing surveillance and full genetic characterization of lagoviruses and other caliciviruses to facilitate detection of future emerging variants of significance to both animal and human health.

## Acknowledgements

We wish to thank all submitters, including domestic pet owners, landholders, and veterinarians, for assistance with sample collection. We thank members of the previous RHD-Boost program of the Invasive Animals Cooperative Research Centre that established the National Rabbit Monitoring Program. We thank Tiffany O’Connor and the virology team at Elizabeth Macarthur Agricultural Institute for contributing additional positive samples. We thank Peter West, Emma Sawyers, and the RabbitScan team for the development and ongoing support of this mobile and web app through which we receive rabbit samples. We thank Roslyn Mourant, Melissa Piper, Dimple Bhatia, and Lily Tran for assistance with sample processing. We would also like to thank Carlo Pacioni for his advice on analyses, and Alex Gofton and Matthew Neave for critical appraisal of the manuscript. Finally, the authors acknowledge the Sydney Informatics Hub and the University of Sydney’s high-performance computing cluster Artemis for providing the high-performance computing resources that contributed to the research results reported within this paper.

## Data availability

Full genome sequences are available in GenBank under accession numbers MW460205 – MW460242. All sequence alignments, tree files, and BEAST xml files are available at https://doi.org/10.25919/758f-4t15.

